# Neural rhythms as priors of speech computations

**DOI:** 10.1101/2025.05.06.652542

**Authors:** Nand Chandravadia, Nabil Imam

## Abstract

The transformation of speech into discrete linguistic representations forms the basis of speech recognition. Natural speech encodes cues at distinct timescales: phonetic features have modulation frequencies of 30-50 Hz, syllables and words around 4-7 Hz, and phrases 1-2 Hz. Strikingly, these frequencies mirror frequencies of endogenous network rhythms of the brain and synaptic time constants of the underlying neural circuits. Here, we suggest that endogenous brain rhythms serve as priors for speech recognition, encoding knowledge of speech structure in the dynamics of network computations. In a network of coupled oscillators, we find that speech is readily identified when characteristic frequencies of the oscillators match frequencies of circuit rhythms in the brain. When signal and circuit rhythms are mismatched, speech identification is impaired. Compared to a baseline recurrent neural network without intrinsic oscillations, the coupled oscillatory network has significantly higher performance in speech recognition across languages, but not in the recognition of signals that lack speech-like structure, such as urban sounds. Our results suggest a central computational role of brain rhythms in speech processing.

## Introduction

Parsing a continuous stream of speech within mixtures of unrelated sounds relies on coordinated neural processes in the brain that track temporal fluctuations of the speech envelope. The auditory cortex displays characteristic patterns of neural activity, notably low-frequency neural oscillations (“rhythms”) in the delta (*δ*) and theta (*θ*) bands, and heightened sensitivity to stimulus modulations in this range [1–3]. Interestingly, temporal modulations of speech envelopes exhibit rhythms in a similar low frequency range, with envelope spectra typically peaking at around 4 Hz [4–6]. When the envelope is artificially compressed, speech comprehension significantly degrades, but recovers when silent gaps are inserted to restore the natural temporal structure of the signal [7]. These observations suggest that the natural frequencies of auditory neural circuits provide a temporal scaffold upon which speech computations take place.

Temporally structured neural activity patterns in the brain tile a range of roughly three orders of magnitude from ultra-slow (< 0.01 Hz) to ultra-fast (∼ 600 Hz) oscillations [8]. Within this broad range are five functional oscillation classes—delta (*δ*), theta (*θ*), alpha (*α*), beta (*β*), and gamma (*γ*)—whose interactions are presumed to subserve various functions, including speech and language processing. It has been proposed that networks of inhibitory and excitatory neurons underlying these oscillations give rise to larger computational units that function as coupled harmonic oscillators [4, 9–11]. These networks of oscillators exhibit periodic (“wavelike”) behavior characterized by frequency, amplitude, and phase, with coupling strengths dynamically adjusted in response to external stimuli. Coupling between oscillators operating at different timescales mirror hierarchical temporal information integration in the brain.

Here, we use model networks of globally coupled oscillators [11, 12] trained with tools from machine learning to investigate endogenous network rhythms of the brain and their relationship to the natural rhythms of input speech signals. The dynamics of each network is defined by the characteristic frequency and damping coefficient of individual oscillators, as well as the their coupling strengths which are learned from speech data. These intrinsic and learned network parameters determine network trajectories over time, which are decoded to categorize speech utterances. We demonstrate that: (1) encoding *a priori* low-frequency brain-like rhythms in the oscillators significantly improve speech recognition performance compared to conventional recurrent neural networks, (2) these computational benefits are consistent across three spoken languages: English, Arabic, and Bengali, (3) network rhythms that align best with the temporal modulations of the speech envelope provide the greatest performance gains across all speech tasks, and (4) these benefits are absent in a non-speech task involving the recognition of urban sounds that lack the temporal structure of speech. These results suggest that low-frequency brain rhythms serve as a dynamic scaffold for speech processing, where coupled oscillations act through the superposition and interference patterns of waves to efficiently extract structured temporal patterns in speech.

## Results

### Rhythms of Speech

The temporal modulations of the speech envelope have a peak energy concentration between 1-5 Hz [4, 13]. These modulations reflect the natural phrasal and syllabic rhythms of spoken language (Fig. 1A). The envelope, which represents the slow amplitude fluctuations of the speech signal over time, encodes the overall intensity pattern of speech and reflects essential rhythmic features of spoken language such as syllabic stress and phrasal timing. The hierarchical organization of language is captured in these low-frequency modulations, with slower components conveying phrase-level structure and faster ones conveying syllabic transitions, enabling the segmentation and interpretation of continuous speech.

**Figure 1:**
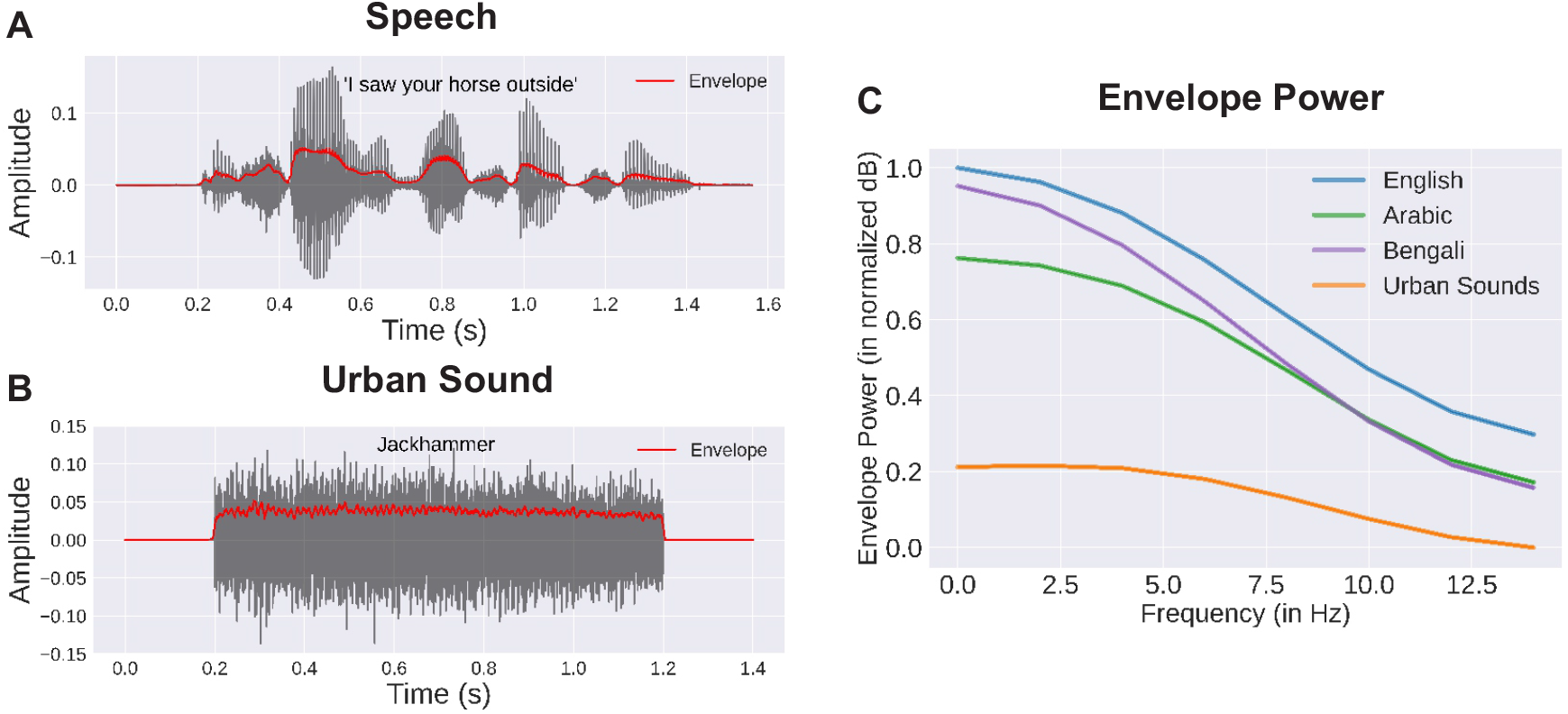
Dynamics of speech and non-speech stimuli. **(A)** Raw audio of the spoken sentence “I saw your horse outside” (grey) with the corresponding speech envelope (red). **(B)** Raw audio of an urban sound (‘Jackhammer’) (grey) and its envelope (red). **(C)** Average power (in normalized dB, [0, 1]) of the envelopes in three spoken language datasets—English, Arabic, and Bengali—and for a dataset of urban sounds.

Unlike the structured and hierarchical rhythms of speech, urban sounds encompass a wide variety of auditory patterns, including irregular noises like car horns or construction activity (Fig. 1B). These sounds lack the temporal regularity and predictable modulations characteristic of speech, instead featuring abrupt onsets, wide spectral distributions, and overlapping frequencies. For instance, while the speech envelope predominantly occupies low-frequency bands between 1-5 Hz, urban sounds span a broader range of frequencies with erratic amplitude fluctuations that lack hierarchical organization. The envelope of urban sounds reflects these irregularities, with peaks and valleys occurring sporadically rather than in systematic rhythmic patterns.

To assess differences in the modulation spectra of speech and urban sounds, we analyze the envelope power in datasets of spoken words across three languages—English, Arabic, and Bengali—and compare it to that of urban sounds encountered in daily life (Fig. 1C). For this, we first compute the amplitude envelope for each sample in the training corpus of each dataset (English, Arabic, Bengali, Urban Sounds). Then, we calculate the power spectrum of each envelope and average them to obtain a representative modulation spectrum. Across languages, we observe a distinctive pattern: the speech envelope peaks between the 1–5 Hz frequency range, reflecting the regular rate at which speakers of different languages convey phonetic information (Fig. 1C, English, Arabic, Bengali). In contrast, urban sounds lack a putative rhythmic pattern, exhibiting no clear peak in the power of the envelope (Fig. 1C, Urban Sounds). The dataset for each language consists of spoken words from different speakers, while the dataset for urban sounds consists of common daily sounds, such as a jackhammer and car horn (Methods).

### Rhythms in Recurrent Neural Networks

The regular temporal structure of speech raises the question of whether the auditory system encodes innate structural knowledge of the signal prior to language acquisition. What computational advantages could this provide? To investigate, we construct a network of coupled oscillators (Fig. 2) to model population-level activity in the auditory cortex [4, 11–14]. Each oscillator is defined by a characteristic frequency (rhythm) and a damping coefficient, which together determine its intrinsic response (Methods). The oscillators are simultaneously driven by this intrinsic dynamic and by raw speech input that modulate network activity. The network adopts an all-to-all coupling architecture, in which the activity of each oscillator influences every other unit, including itself. Internal couplings—determining how oscillators influence each other’s activity—are initialized randomly, but adapt through training to encode stimulus structure. As a result, the network learns a rich and diverse set of trajectories in its state space, shaped by its intrinsic rhythms and the temporal regularities of speech.

**Figure 2:**
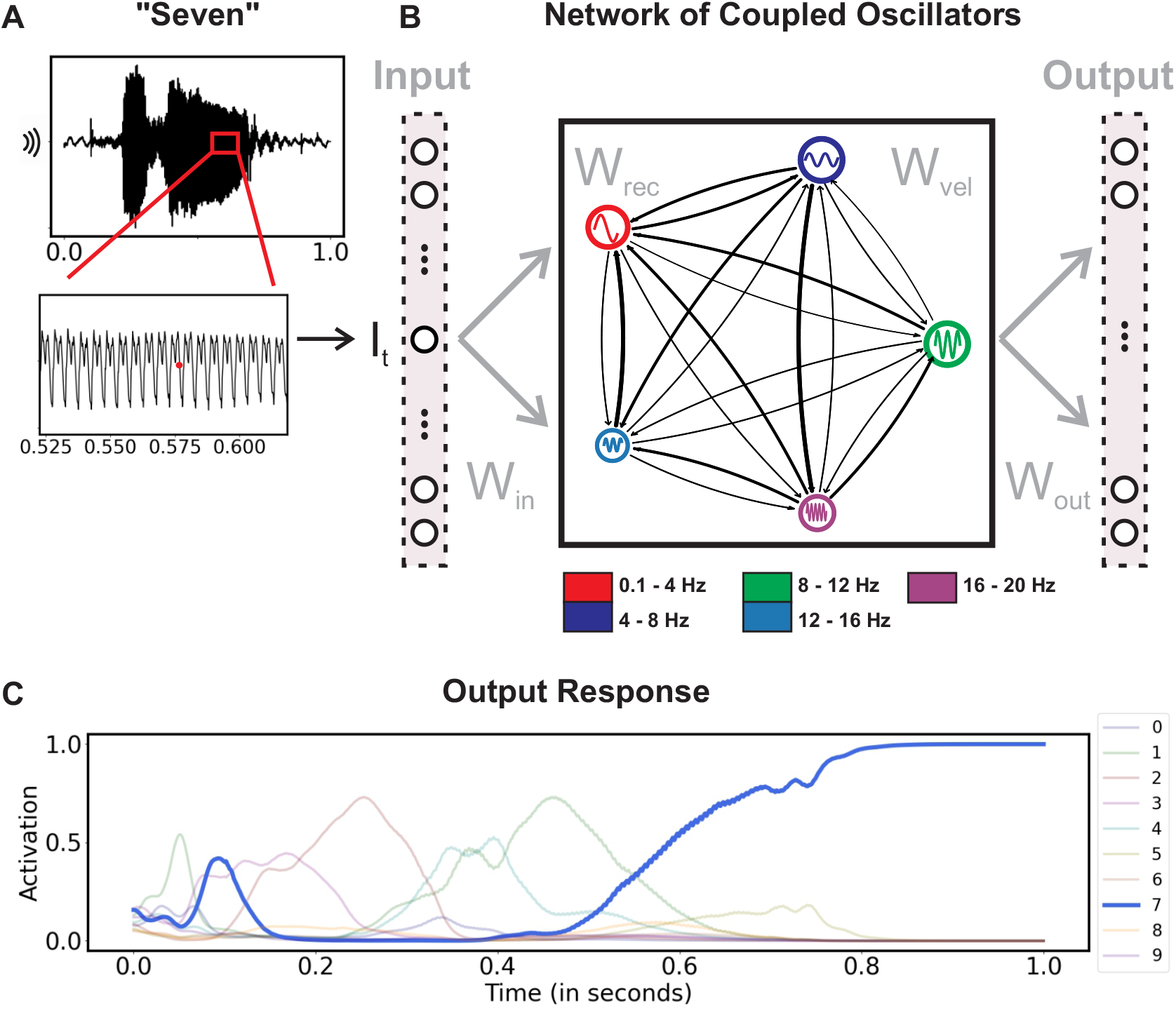
Network architecture. **(A)** Time-varying speech waveform of the spoken word “seven,” sampled at each time point *I*_*t*_. **(B)** Illustration of a network of coupled oscillators receiving samples of the speech waveform, with plastic input (*W*_*in*_), recurrent (*W*_*rec*_, *W*_*vel*_), and output (*W*_*out*_) weights. **(C)** Activity in the output layer in response to the spoken word.

We configure the oscillators with heterogeneous frequencies drawn from a fixed range. We evaluate several frequency ranges: some corresponding to brain oscillations (e.g., delta, theta, and gamma bands), while others having little correspondence to brain activity. During learning, we present raw auditory waveforms to the network, and the network adapts its coupling parameters such that the waveforms—either speech or non-speech—are classified into appropriate categories (Methods). For instance, repeated presentations of the spoken word ‘seven’ guide the network toward a dynamical trajectory from which a linear readout layer determines the stimulus class. Once trained, we present the network with unseen samples of the learned categories, and record its recognition performance. The dynamics of the raw waveform we utilize here is reflected in its time–frequency spectrum, though the waveform itself offers a finer temporal resolution.

### Performance Across Languages

To assess the effects of endogenous network oscillations on speech processing, we evaluate the performance of our model network in a series of tasks, including spoken word recognition in three different languages, and recognition of altered speech and non-speech sounds.

First, we initialize a network of 256 oscillators, with characteristic frequencies drawn uniformly at random from the range 0.1 Hz to 20 Hz. We train this network on a dataset of English spoken words, consisting of utterances of the words “zero” through “nine” by different speakers. Each spoken word in the training set was uttered multiple times. During training, we feed the network the raw speech waveform (as recorded from the speaker), and adjust the coupling strengths via the back-propagation through time (BPTT) learning algorithm (Methods) [15, 16]. After training the network for several epochs, we evaluate its word recognition performance on a held-out test set. As shown in Fig. 3A, the network achieves robust accuracy within the first few epochs, reaching a final performance of 89% ± 1% after 50 epochs. Accuracy values are averaged over 10 random initializations of network parameters. Notably, this high test performance is achieved using only the raw speech waveform as input—without any pre-processing or feature extraction—during both training and testing.

**Figure 3:**
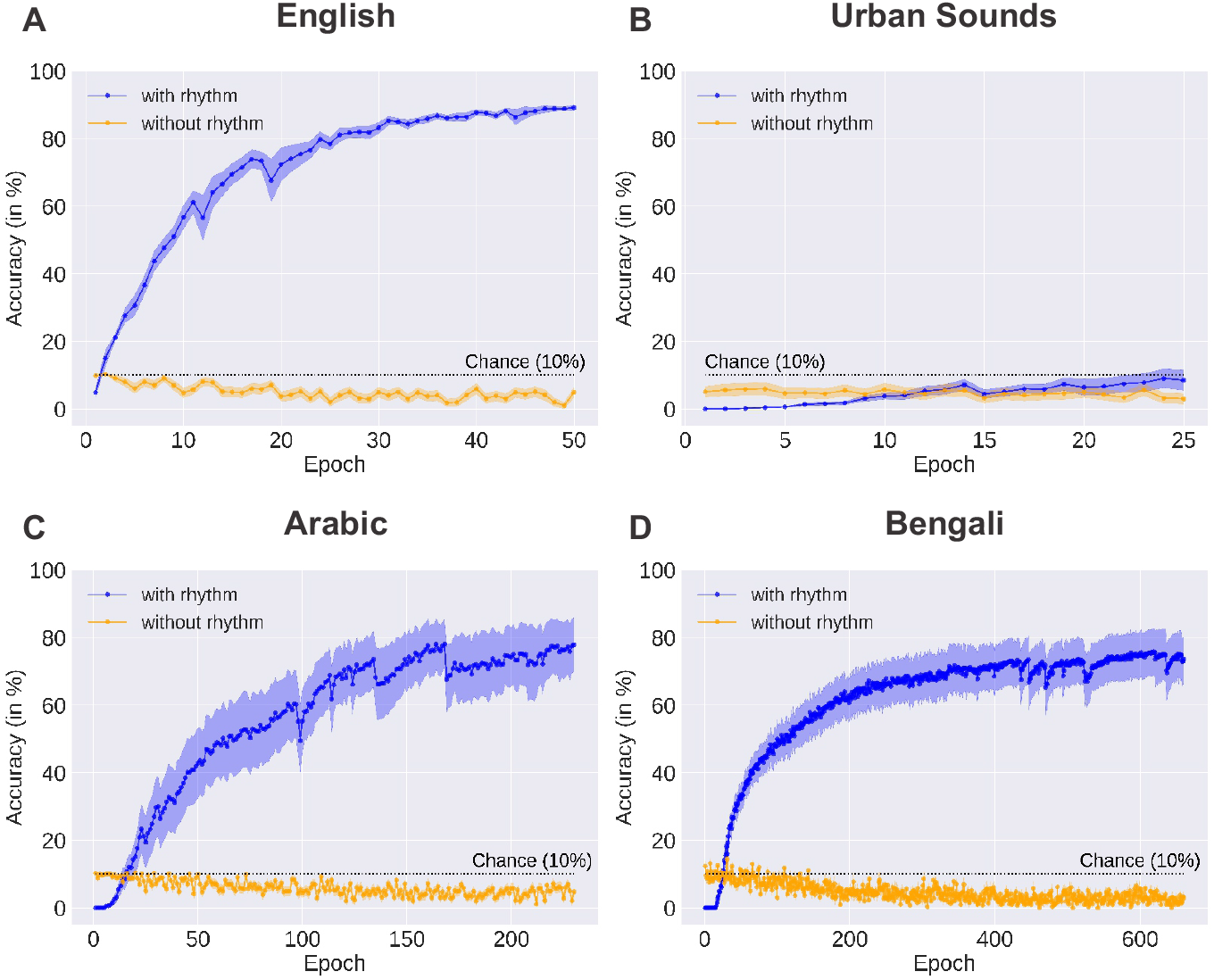
Performance with and without network oscillations. Test accuracy of networks configured with oscillation frequencies chosen uniformly at random from the range [0.1 Hz, 20 Hz] (blue), compared to the same networks without oscillations (yellow). The accuracy after every training epoch is shown. Values are averaged across ten random seeds. Shaded region depicts standard error. Datasets were **(A)** AudioMNIST (English), **(B)** UrbanSound8k (Urban Sounds), **(C)** Arabic Spoken Digits, **(D)** Bengali Spoken Digits.

For comparison, we train a recurrent neural network without endogenous oscillations on the same dataset of spoken English digits, using the same learning algorithm and raw speech input (Methods). Unlike the oscillatory network, this recurrent neural network was unable to learn the task (Fig. 3A). Test accuracy remained below chance level at 5% ± 2% throughout training.

Next, we assess which oscillator frequencies are most effective in recognizing speech. We initialize different networks, each configured with frequencies drawn uniformly at random from a specific range. We find that networks with oscillator frequencies in the range 1-5 Hz achieve the highest accuracy and training efficiency (Fig. 4A). Interestingly, this corresponds to the peak envelope power of the speech signal (Fig. 1C), indicating that a *match* between signal and network oscillations leads to the best accuracy. Accuracy dropped steeply for higher frequency network oscillations, falling to chance levels for frequencies above 40 Hz.

**Figure 4:**
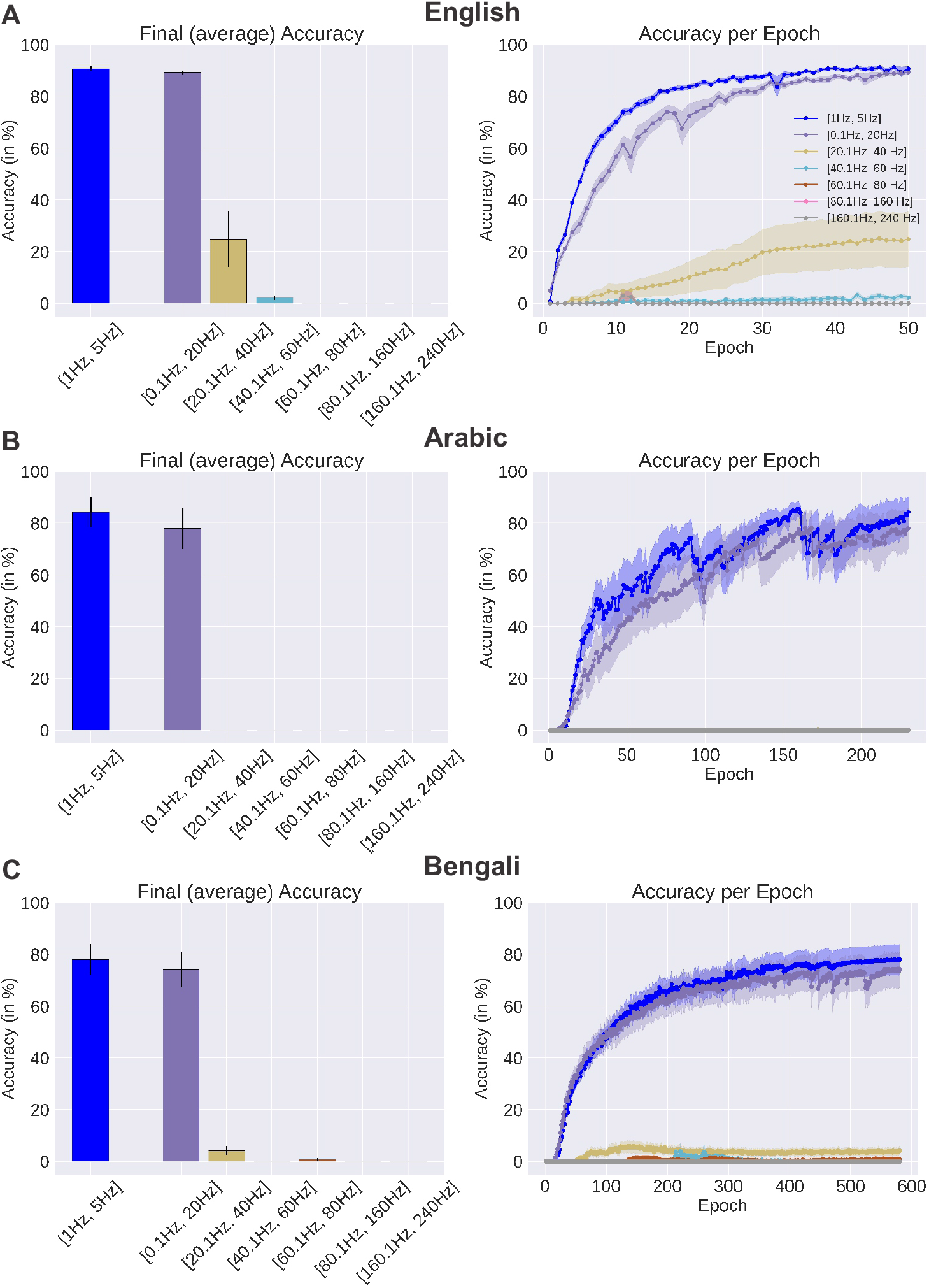
Performance across oscillation frequencies. **(A)** Accuracy in detecting English spoken words. Left: Test accuracy of networks configured with oscillator frequencies drawn uniformly at random from the specified ranges. Values are averaged across ten random seeds. Error bars depict standard error. Performance is highest for low-frequency oscillations that match the temporal structure of the signal. Right: Test accuracy per epoch of the same networks. **(B, C)** Arabic, Bengali

These results indicate that the coupled oscillatory network achieves high performance when it encodes knowledge of the temporal structure of speech—specifically, its amplitude modulation frequencies—prior to training. To test whether this effect extends beyond English, we evaluate the network on two additional datasets: Arabic and Bengali spoken digits.

The Arabic and Bengali spoken digits datasets contain utterances of the digits “zero” through “nine” in Arabic and Bengali respectively, each spoken multiple times by different speakers. As before, the network is trained using the raw speech waveform without any pre-processing or alignment. The results closely mirror those observed for English. Networks configured with low-frequency oscillations achieve substantially higher performance (78% ± 8% for Arabic, 74% ± 7% for Bengali) compared to networks that lacked endogenous oscillations (5% ± 2% for Arabic, 3% ± 1% for Bengali, both below chance; Fig. 3C-D). Networks configured with higher-frequency oscillations (*>* 20 Hz) are unable to learn, with performance dropping to chance levels (Fig. 4B-C), again showing that a match between signal and network oscillations is necessary for high task performance.

To examine which features of the spoken words are most critical for learning, we decompose the speech waveform into mel-frequency cepstral coefficients (MFCCs), which extracts frame-by-frame spectral content from continuous audio signals. The first MFCC coefficient computes the overall log-energy of the signal and closely tracks the speech envelope (Fig. S1B), while higher-order coefficients capture progressively finer spectral details (Fig. S1C). When the first MFCC is provided as input to the high-performing oscillatory networks, they successfully learn the task (Fig. S2B). However, this did not hold for any of the other coefficients; for example, when the same networks are given the seventh MFCC, performance remains at chance level (Fig. S2C). This indicates that the oscillatory networks preferentially extract envelope-level information from the raw waveform. They achieve high performance by tracking the temporal modulations of the envelope, which is the most salient structure in the signal. Networks without endogenous oscillations, which are unable to learn the task using the raw waveform, achieve high performance when given the first MFCC, since the MFCC transform extracts envelope information by design (Fig. S2B).

### Performance for Non-Speech Sounds

Next, we examine whether the results for speech generalize to non-speech stimuli using a dataset of urban sounds [17]. Urban sounds, such as traffic or construction noise, lack the distinct temporal modulations characteristic of speech. We test whether networks configured with low-frequency oscillations still provide a performance advantage when trained to classify sounds in this dataset. The results indicate that these networks are no more effective than non-oscillatory networks in this task. Performance values are 8% ± 3% for networks with oscillations and 2% ± 2% for networks without oscillations, both below chance level (Fig. 3B). This provides further support for our hypothesis that endogenous oscillations serve as prior knowledge of signal structure—the priors that are effective at speech recognition are entirely ineffective in recognizing signals that lack speech-like structure.

Speech rhythms can vary naturally across speakers and contexts, prompting the question of how such variability affects models with fixed temporal assumptions. In particular, we asked whether artificially altering the speech rate—by slowing it down (0.5 ×) or speeding it up (2 ×)—impacts our highest performing networks. The results show a significant drop in performance of ∼ 20% when we either speed up or slow down the speech rate, compared to the natural speech rate (Fig. S3). The observed performance drop reflects a misalignment between the network’s intrinsic expectations and the altered speech rates.

To further examine signal and network alignment, we train networks on synthetic sinusoidal inputs of fixed frequencies. The network oscillators are initialized with frequencies drawn from fixed ranges and their task is to classify the frequency of the input signal. This setup allows us to directly test whether matching signal and network rhythms facilitate task learning. When the task is to classify sinusoids drawn from the frequency range 10-100 Hz (requiring the signal to be classified into one of 10 uniformly-spaced frequency bins), a network with oscillator frequencies drawn from the same range learns rapidly, achieving nearly 99% accuracy (Fig. 5). In contrast, a network with oscillator frequencies drawn from the range 1-10 Hz performs at chance level, while a network with frequencies drawn from the range 100-110 Hz learns the task slowly, plateauing to around 80% accuracy. As a control, a network without intrinsic oscillations also learns the task slowly and plateaus at 80%.

**Figure 5:**
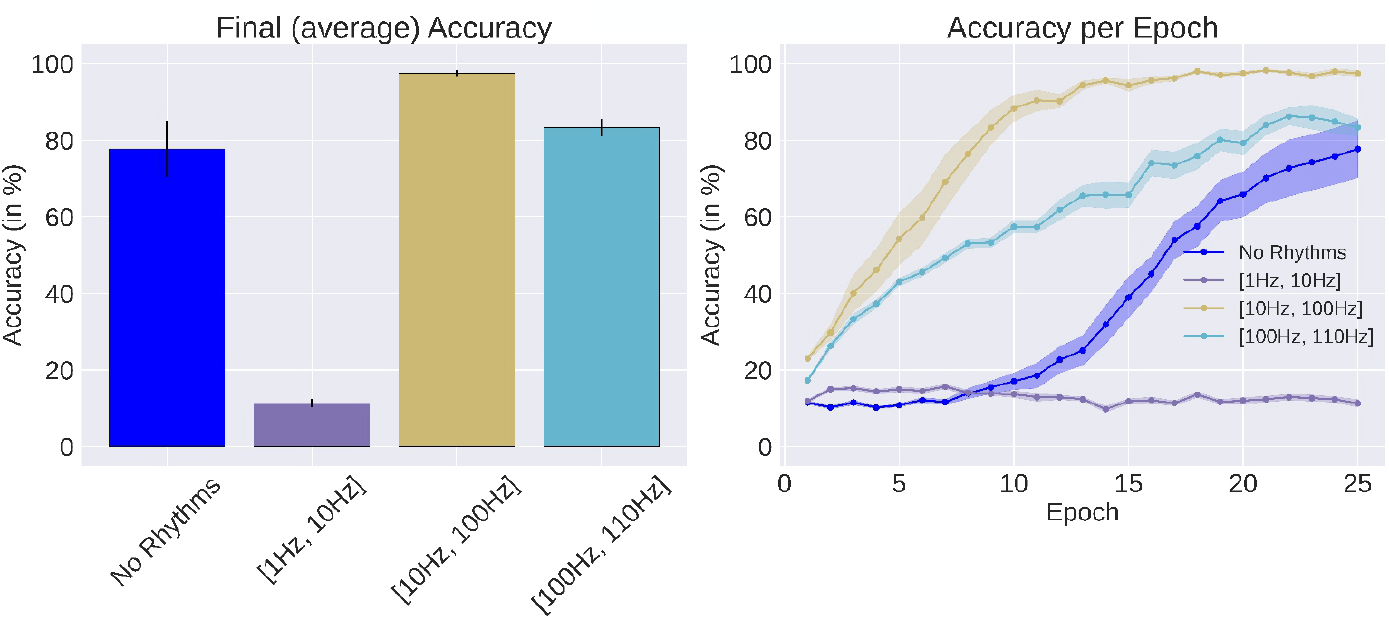
Performance in classifying synthetic sinusoidal inputs drawn from the range 10-100 Hz. Left: Test accuracy of networks configured with oscillator frequencies drawn uniformly at random from the specified ranges. Values are averaged across ten random seeds. Error bars depict standard error. Networks with matching intrinsic frequencies (10–100 Hz) achieve best performance. Right: Test accuracy per epoch of the same networks.

Importantly, networks that achieve high performance in this sinusoid recognition task exhibit learned rhythmic activity whose period matches the period of the input signal, whereas networks performing at chance do not show this signal-network alignment (Fig. S4). These results reaffirm that a match between signal and network rhythms is essential for achieving high task performance. They also indicate that while networks with low-frequency endogenous oscillations are unable to detect features of faster inputs, those with high frequency retain the ability to adapt downward to slower inputs in this particular task.

### Effects of Damping

Each node in our network is parameterized by a fixed frequency coefficient *γ* and fixed damping coefficient *ϵ* (Methods). The choices of these coefficients bias the learning process toward particular oscillations. In the results presented so far, we have examined the effects of the frequency coefficient while keeping the damping unchanged across the setups. In this section, we vary the damping coefficient and assess its effects on task performance.

First, in the English spoken words task, we set the frequency coefficient to zero and choose damping values for each unit uniformly at random from the range [0.1, 80]. We observe that this network initially struggles with the task but gradually reaches high performance with continued training (Fig. S5). High performance is achieved once the network learns to oscillate at the frequency of the speech envelope (∼ 1 Hz, Fig. S6). In contrast to this slow learning, if the same network is initialized with frequency coefficients matched to the frequency of the speech envelope, it requires fewer training epochs to reach high performance, learning the task more rapidly (Fig. S5).

Next, we perform a systematic scan across a wide range of damping coefficients and assess performance. For this, networks are initialized with frequencies that match the speech envelope, while the damping range is varied across intervals of progressively increasing range. We find that when damping is too low (or absent), networks perform poorly (Fig. S7A); for example, test accuracy is 5% when the damping is chosen from the range [0.1, 5] uniformly at random for each unit. Performance improves once damping is increased (e.g., to the range [0.1, 45]) but when damping becomes too large, performance drops again (Fig S8). These results indicate that there is a sweet spot for damping: too little damping prevents network oscillations from stabilizing to the task, while too much damping suppresses oscillations altogether (Fig. S9).

### State-Space Dynamics

To visualize the dynamics of the networks, we plot the first three principal components of network activity in response to spoken words, using network configurations that achieved best task performance. We find that before training, network trajectories across words and languages are almost identical, with only minor variations reflecting subtleties in intonation and stress patterns (Fig. 6A-C: Untrained). During training, the trajectories become increasingly distinct for each spoken word, and once training is complete, the trajectories are highly decorrelated, encoding the specific patterns of each word across the three languages (Fig. 6A-C: Trained). In contrast, networks that lack intrinsic oscillations do not learn decorrelated representations across words and languages (Fig. 7). The scaffold provided by the low-frequency oscillations, therefore, enable class-wise trajectory separation via training, which is the reason underlying high classification accuracy (Fig. 3).

**Figure 6:**
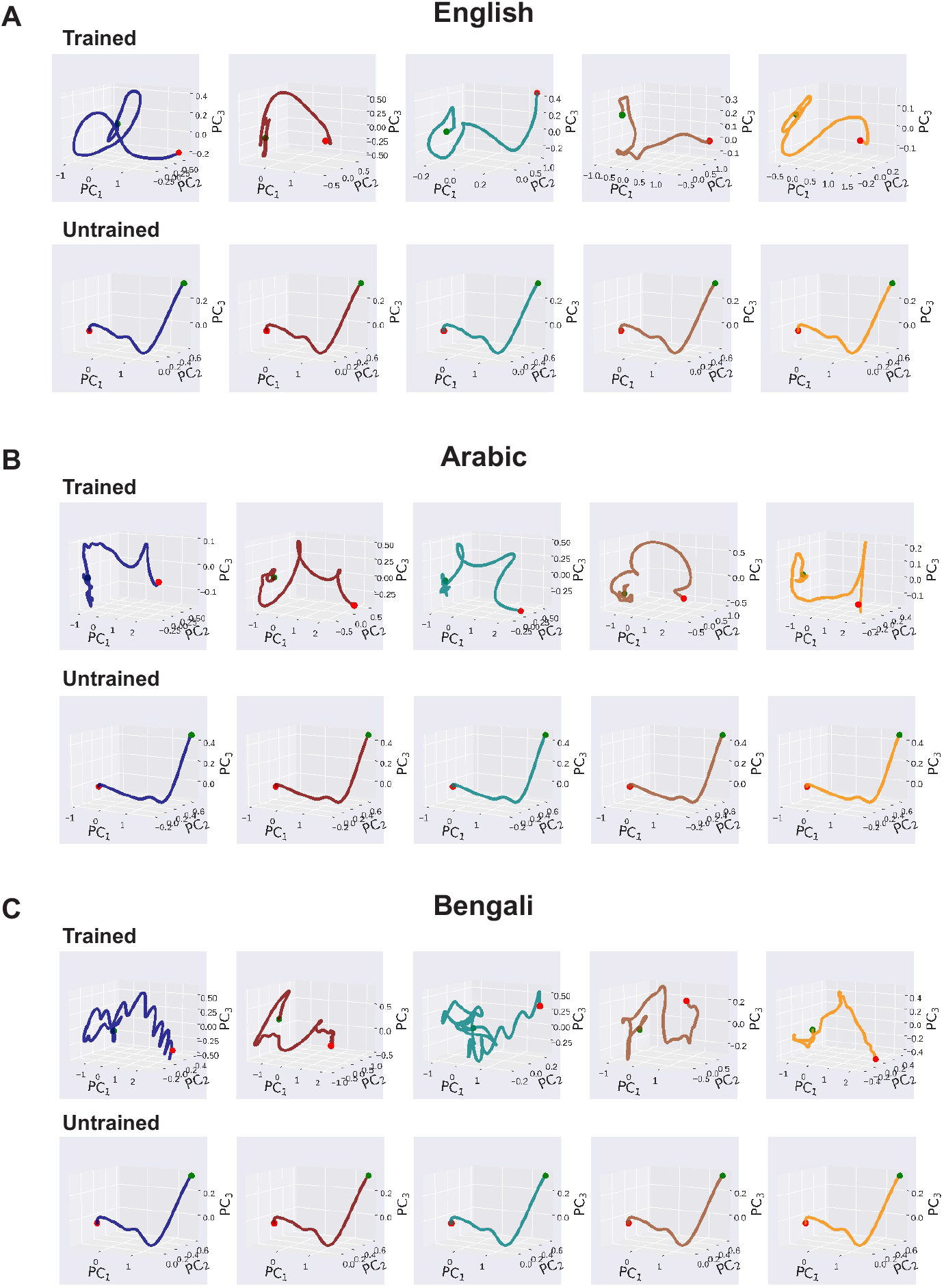
Network trajectories with endogenous oscillations. **(A)** Trajectory of network activity in PCA space in response to five spoken digits in English (“zero”, “two”, “four”, “six”, “eight”). Trajectories are averaged across all test samples in the dataset. The network is initialized with frequencies drawn uniformly at random from the range [0.1 Hz, 20 Hz]. Green and red dots depict start and end of the trajectories respectively. Top: before training. Bottom: after training. **(B, C)** Arabic, Bengali.

**Figure 7:**
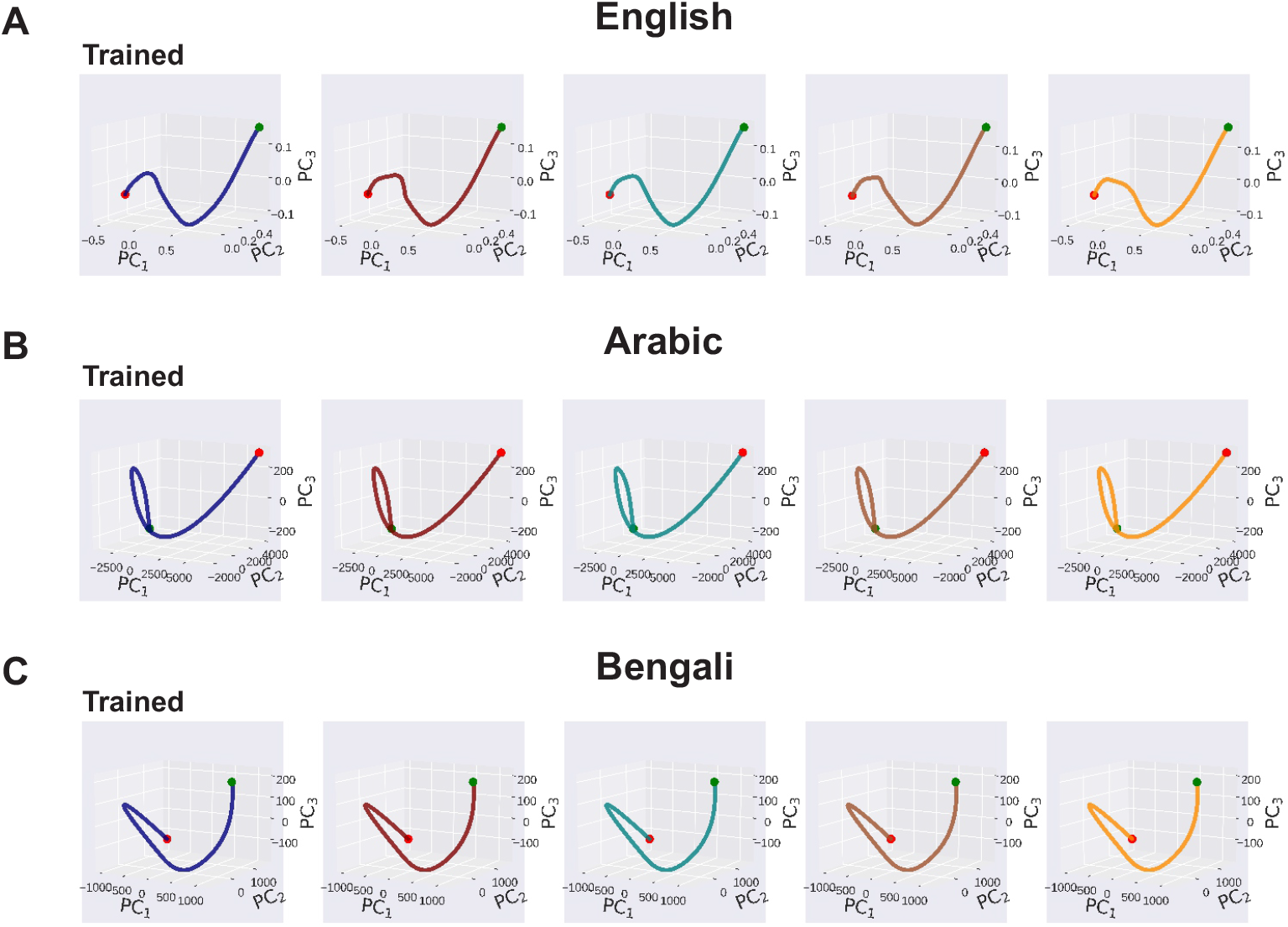
Network trajectories without endogenous oscillations. **(A)** Trajectory of network activity in PCA space in response to five spoken digits in English (“zero”, “two”, “four”, “six”, “eight”) for a network with no endogenous oscillations and randomly initialized weights. Different digits evoke the same trajectory. Trajectories are averaged across all test samples in the dataset. Green and red dots depict start and end of the trajectories respectively. **(B, C)** Arabic, Bengali.

We then assess which oscillators predominantly shape activity when a network is initialized with frequencies drawn uniformly at random from the range [0.1 Hz, 20 Hz]. For this, we calculate the component loadings of the first principal component of the network’s activity after training. The first principle component captures ∼ 70% of the total variance. Across the three languages, we observe that oscillators within 0.1 Hz and 2.5 Hz dominate the network’s activity (Fig. 8), with higher-frequency oscillators (*>* 5 Hz) exerting only a weak influence. Thus, consistent with our other results, low-frequency oscillations are the primary drivers of computation in the network.

**Figure 8:**
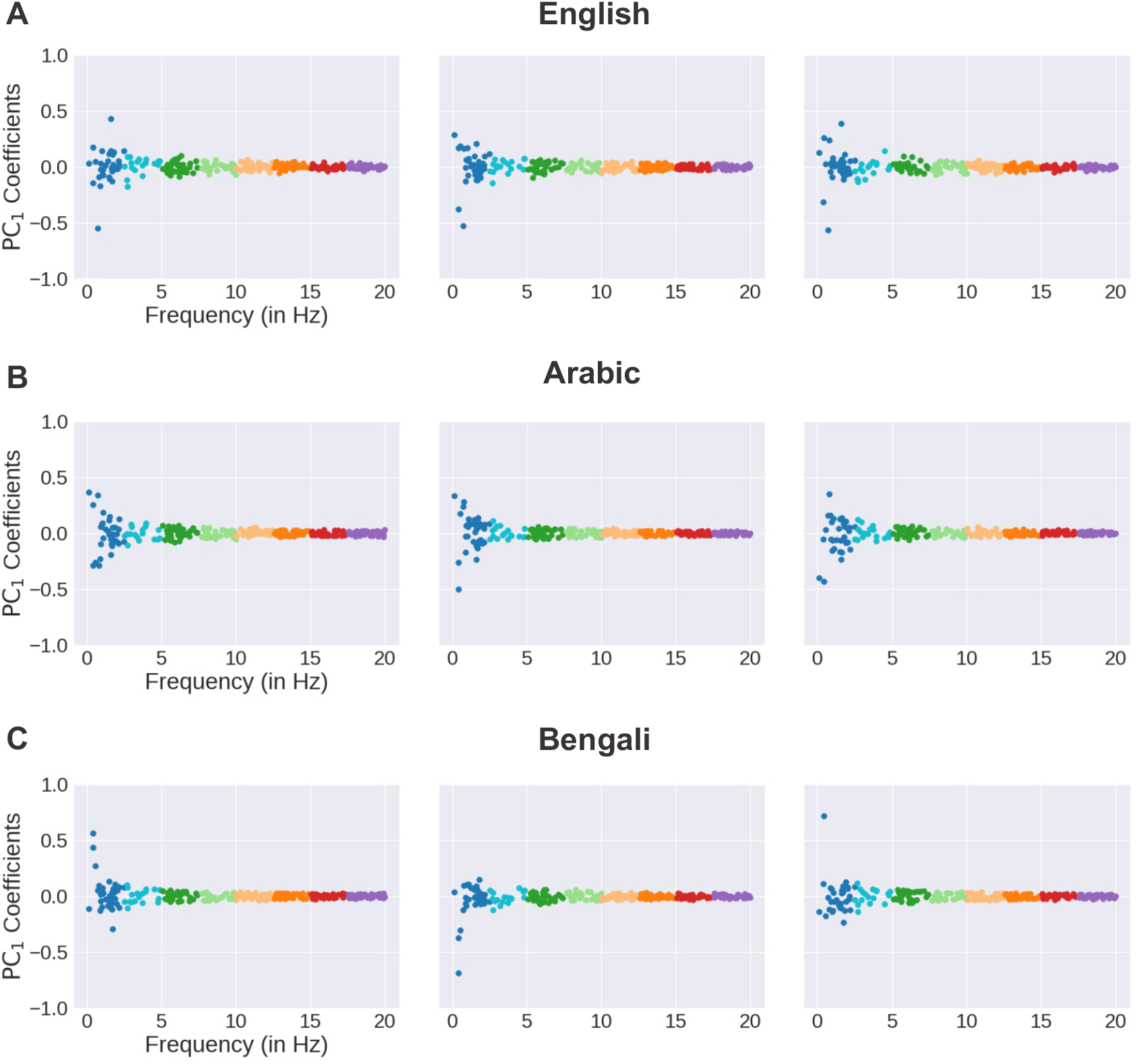
Dominant oscillation mode. The influence of each oscillator to the first principal component in response to **(A)** English, **(B)** Arabic, and **(C)** Bengali spoken words for a network trained with randomly initialized frequencies from the range [0.1 Hz, 20 Hz].

## Discussion

We show that networks of coupled oscillators with endogenous low-frequency oscillations accurately learn to recognize spoken words in three distinct languages. Our results demonstrate that oscillations serve as prior knowledge of speech structure, providing a temporal scaffold on which statistical learning takes place. Unit oscillations drawn from the 1-5 Hz frequency range results in the best task performance in all three languages. This suggests that despite differences in phonetic and prosodic features of English, Arabic, and Bengali, their raw acoustic signals share common, perhaps universal, temporal structure. It has been proposed that speech production involves quasi-rhythmic motor patterns dictated by skeletal, muscular, and neural circuit-level constraints of the human speech-motor system [4, 18, 19]. The perceptual system, in turn, imposes rhythms that are tuned to detect and interpret speech signals at the appropriate temporal granularity, akin to active sensing in other modalities, but without explicit movement-driven sampling. Our results lend computational support to this view.

Imposing oscillatory dynamics in itself is insufficient for extracting task relevant structure. Rather, a match between signal and network oscillations is essential. We find that networks with intrinsic oscillations mirroring those of the audio input reach high speech recognition accuracy, whereas networks with oscillations mismatched to the signal are unable to learn the task. In circuit theory, frequency matching between system and input, or *resonance*, imparts frequency selectivity, voltage magnification, and other signal transformations, widely used in analog communication systems. In analogy, circuit resonance in the brain has been reported in several studies, where signals arriving in phase with the natural oscillations of cortical circuits are selectively gated or amplified [20–22]. In our model, we use an explicit parameter *γ* to define the characteristic frequency of each node, and suggest it be interpreted as a *dynamic* prior that enables effective categorization of signals with matching temporal structure. We note that although the initial resonance properties of each unit is determined by *γ*, the learned frequency response is more complex due to network influences, as also reported in other studies [11, 12].

In machine learning, priors, or inductive biases, refer to pre-existing knowledge or initial assumptions given to a model as cues to guide the learning process. This enables faster and more efficient training. A prior may be encoded by imposing a particular structure on the form of a model or as a hint about the underlying distribution of the input. Whether explicit or implicit, priors lend direction to the learning algorithm to find meaningful patterns in the data, narrowing the range of possible hypotheses about the class of functions being learned [23]. In a neural network, finding good minima in a high-dimensional, non-convex loss landscape can be difficult [24]. Priors help circumvent this difficulty by initializing the network to favorable configurations, enabling it to avoid adverse terrain of the loss landscape.

In the brain, priors come encoded at birth, though their precise forms are not well understood. They may take the form of stereotyped circuit architectures imposed by development [25–28], or as we propose here, particular types of dynamical response. Learning involves adapting these initial expectations to the patterns of stimuli encountered in the world. For example, in language acquisition, priors may encode general syntactic constraints, with exposure to a particular language resulting in language-specific expectations and associations with sound. The learning rules in the brain likely differ substantially from the BPTT algorithm used here [29–32], although alternative forms of credit assignment may be partly at play [33–35].

Network structure in the brain has inspired inductive biases in artificial neural networks [36–38]. The topographic organization of the primary visual cortex, for example, where neurons respond selectively to localized regions of the visual field, has motivated the use of local filters in convolutional neural networks. By enforcing spatial locality and translational invariance, these architectural choices encode knowledge essential for efficient processing of visual information [39, 40]. Beyond this spatial organization, neural systems exhibit rich temporal dynamics, often exhibiting traveling waves of activity patterns that unfold across the cortical surface. When embedded within recurrent neural networks (RNNs), traveling waves generate dynamic representations that can be exploited for efficient sequence learning, continual learning, and long-term memory consolidation [41–46]. Other phenomena characteristic of coupled neural systems, including transient synchronization, phase locking, and the hierarchical nesting of oscillations, may also be incorporated into similar wave-based models. These wave-RNNs offer new tools to artificial intelligence and serve as a platform for testing computational hypotheses about brain function.

## Methods

### Datasets

We used the AudioMNIST (English), Arabic Spoken Digits, BanglaNum (Bengali), and UrbanSound8K datasets in our tasks. AudioMNIST consists of 30,000 audio recordings (∼ 8.3 hours) of spoken digits (0 through 9) across 60 different English speakers [47]. Each recording is ∼ 1 second, originally sampled at 48 kHz on a single-channel. Each speaker recited each digit 50 times, resulting in 3,000 recordings across each class.

The Arabic Spoken Digits dataset is a subset of the Arabic Speech Corpus for Isolated Words [48]. The dataset consists of approximately 5, 000 audio recordings of spoken digits (0 through 9) across 20 different Arabic speakers. Each recording is ∼ 1 second, originally sampled at 44, 100 Hz on two channels (i.e., stereo). The Bengali Spoken Digits (BanglaNum) dataset consists of approximately 2, 250 audio recordings of spoken digits (0 through 9) across 40 different Bengali speakers [49]. Each recording is ∼ 0.5 seconds, originally sampled at 16 kHz on a standard single-channel microphone.

UrbanSound8K consists of field recordings of various sounds found in an urban environment, spanning 10 low-level classes (i.e., air conditioner, car horn, children playing, dog bark, drilling, engine idling, gun shot, jackhammer, siren, and street music) [17]. In total, there are 8,732 audio recordings (∼ 7.06 hours), each with a variable time length (≤ 4 seconds) with sampling rates varying from 8 kHz to 96 kHz.

For the language datasets, we randomly select 90% of the samples to be used as the training set, and the rest as the test set. For Urban Sounds, we randomly select 80% as the training set, and the rest as the test set. For all datasets, we first down sample each recording to 8 kHz, and then normalize the amplitude of the signal to the interval [ − 1, 1]. The resulting time-series, *I*(*t*), is used as the input signal to the network.

For the mel-frequency experiments, the raw waveform was transformed into mel-frequency cepstral coefficients (MFCCs) before training. Each signal was processed with an MFCC transform that extracted 13 coefficients per frame, using a 200-point short-time Fourier transform (STFT) with a hop length of 5 samples and a 20-channel mel filterbank. The resulting time–frequency representation was used as the network input.

For the sinusoidal experiments, we generated synthetic waveforms of fixed frequencies. The waveform was defined as *s*(*t*) = *A* sin(2*πft* + *ϕ*), where *A* is the amplitude, *f* is the frequency, *t* is time, and *ϕ* is the phase shift. For each frequency, we created multiple examples by sampling the amplitude uniformly from [0.1, 5] and the phase shift uniformly from [0, 2*π*]. Using this procedure, we generated 10 classes of sinusoids corresponding to fixed frequencies of 10 Hz through 100 Hz, equally spaced. An analogous dataset was also generated for the low-frequency range [1, 10] Hz, with classes defined by fixed frequencies of 1 through 10 Hz. Each class consisted of 1000 samples. We randomly selected 80% of the samples to be used as the training set, and the rest as the test set. These waveforms then served as the input to the network.

### Model Description

We use the coRNN (Coupled Oscillatory Recurrent Neural Network) setup, which is an RNN architecture of time-discretized coupled oscillators [12]. In the coRNN, the coupled oscillators take the form of the following second-order, *m*-dimensional system of ODEs:

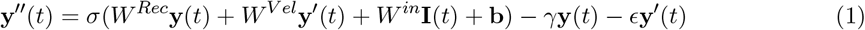

where,

**I**(*t*) ∈ ℝ^*d*^ time-varying input signal

*W* ^in^ ∈ ℝ^*m×d*^ input weight matrix

**y**(*t*) ∈ ℝ^*m*^ hidden states of the RNN

**y**^*′*^(*t*) ∈ ℝ^*m*^ velocity of hidden states of the RNN

*W* ^Rec^, *W* ^Vel^ ∈ ℝ^*m×m*^ recurrent weight matrices

**b** ∈ ℝ^*m*^ bias term

*σ*(·) ∈ ℝ activation function (tanh)

***γ*** ∈ ℝ^*m*^ characteristic frequency (Hz) of each node (≥ 0)

***ϵ*** ∈ ℝ^*m*^ damping of each node (≥ 0)

Each node in the network is parameterized by a fixed characteristic frequency (*γ*) and damping (*ϵ*) coefficient. Time, *t* ∈ [0, 1], represents a continuous variable. To simulate Eq. 1, we let the velocity term, **y**^*′*^(*t*) = **z**(*t*). We can then rewrite the second-order ODE (Eq. 1) as a first-order ODE (Eq. 2), where **z**^*′*^(*t*) = **y**^*′′*^(*t*), resulting in:

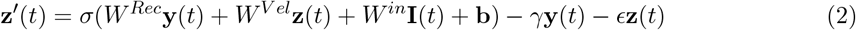

For a fixed timestep, 0 < Δ*t* < 1, the IMEX (implicit-explicit) discretization of this equation results in:

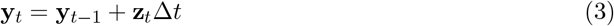

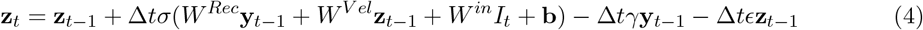

We consider only the explicit discretization of equations 3 and 4 in the network simulations.

We simulate the coRNN both with and without endogenous frequencies, using a network of 256 units in each case. In the endogenous case, we configure the network with various combinations of oscillations and damping (Table S1 in Supplementary Information). To configure networks without endogenous oscillations, we set *γ* and *ϵ* to 0 in Eq. 1, effectively resulting in an Elman (or standard) RNN [37]. All networks have an equivalent number of trainable parameters.

### Network Training

All simulated networks consist of a trainable input, recurrent, and readout (SoftMax) layer, where the initial weights are sampled from a uniform distribution, 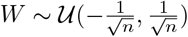 (*n* is the input dimension to each layer). The input to each network, *I*(*t*), is a time-varying signal. All parameters were subsequently trained via BPTT (Backpropagation Through Time). We used the cross-entropy loss function with the AdamW optimizer, a learning rate of 4.2 · 10^−3^, and a time step of 5 · 10^−3^.

For the AudioMNIST, Arabic Spoken Digits, Bengali Spoken Digits dataset, we used 50 training epochs with a batch size of 64, which took a total of ∼ 50 hours of training time per network on a NVIDIA V100 GPU. For the UrbanSounds8k dataset, we used ∼ 25 epochs with a batch size of 32, which took ∼ 25 hours to train. We carried out a systematic scan of the frequency and damping ranges (Table S1 in Supplementary Information), but did not perform an exhaustive grid-search due to the long training times.

### Network Readout

To assess network performance on the speech tasks, we used the following classification scheme: for the last ∼ 62.5 ms of the input signal, we take the node in the SoftMax layer with the highest activation. Specifically, if the maximum SoftMax activation at time *t* exceed a threshold of 0.60, the corresponding class *i* is assigned as the prediction at that time point; otherwise, the output is labeled as Null. After collecting predictions across the classification window, we determined the final class label as follows: if class *i* is the most frequently predicted label in this window and appears in at least 80% of the non-null predictions, then *i* is selected as the network’s final decision. Otherwise, the classification is marked as uncertain. This scheme ensures a robust classification at readout, despite the oscillatory nature of the network. See Fig. 2C for an illustration of a network response.

For the sinusoidal classification task, we adopted a complementary readout strategy better suited to fixed-frequency sinusoids. Rather than using only the final segment of the input, we concatenated the hidden-state activity of all network units across the entire temporal window into a single vector for each trial. A linear readout layer (softmax classifier) was then trained on this aggregated representation to predict the frequency class. This approach gave the classifier access to the entire trajectory of network dynamics, allowing it to capture the oscillatory patterns that persisted throughout the input.

## Supplementary Information

**Table S1:**
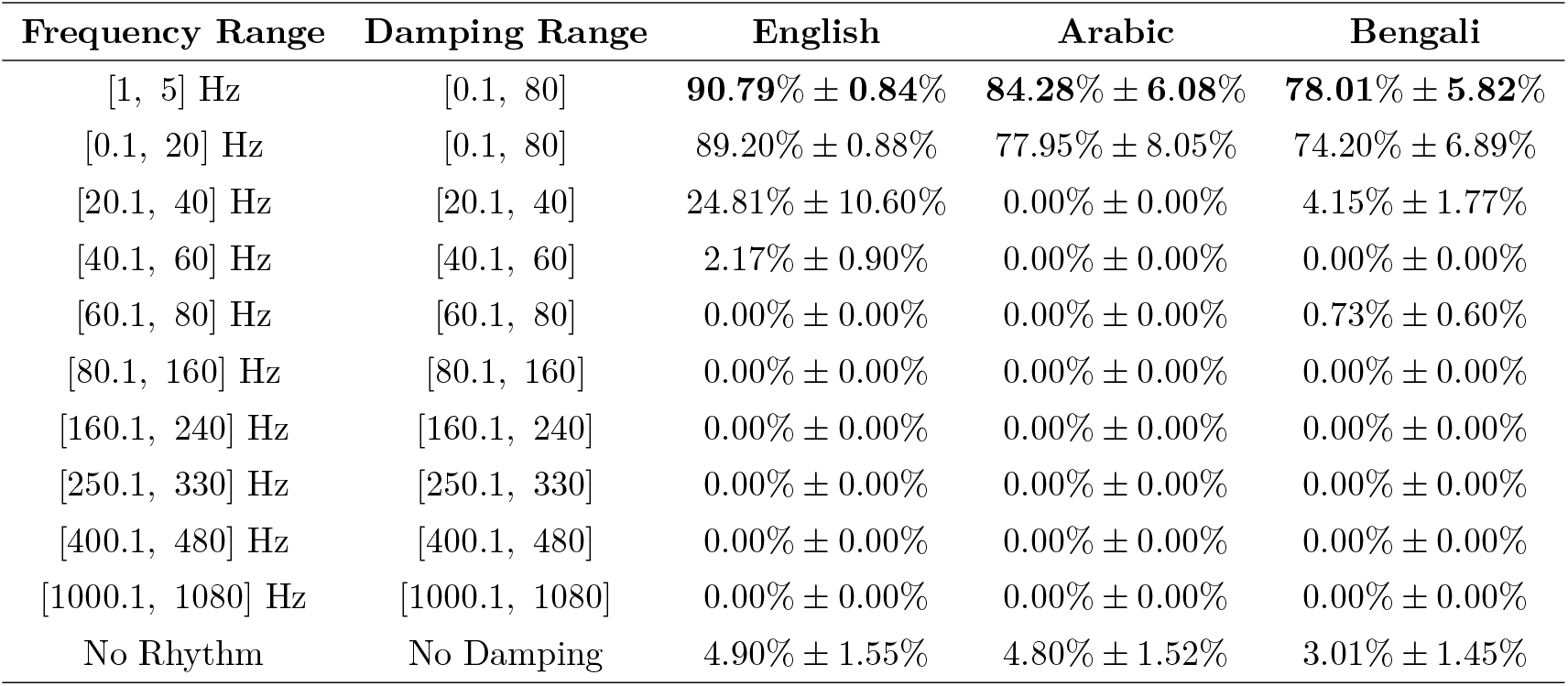
Final test accuracy of networks configured with frequency and damping coefficients selected uniformly at random from the specified ranges. The bold values indicate the highest accuracies. Values are averaged (± standard error) across 10 random seeds.

**Figure S1:**
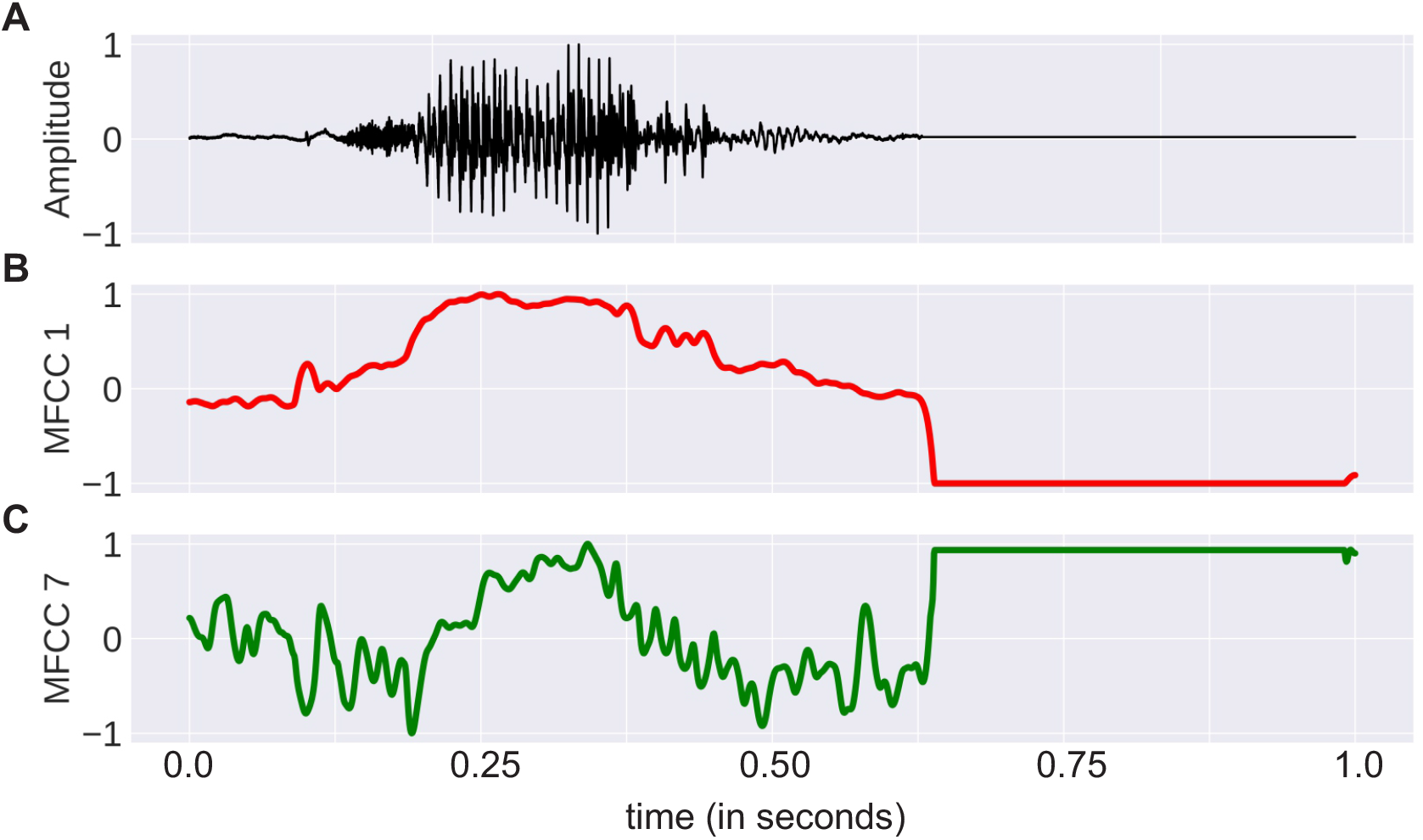
Mel-frequency cepstral coefficients (MFCCs) extracted from raw speech waveform. **(A)** Waveform of the spoken word “zero.” **(B)** First MFCC coefficient, which closely resembles the speech envelope. **(C)** Seventh MFCC coefficient, which reflects finer spectral details. All MFCCs were normalized to [-1, 1] for comparison with the raw waveform.

**Figure S2:**
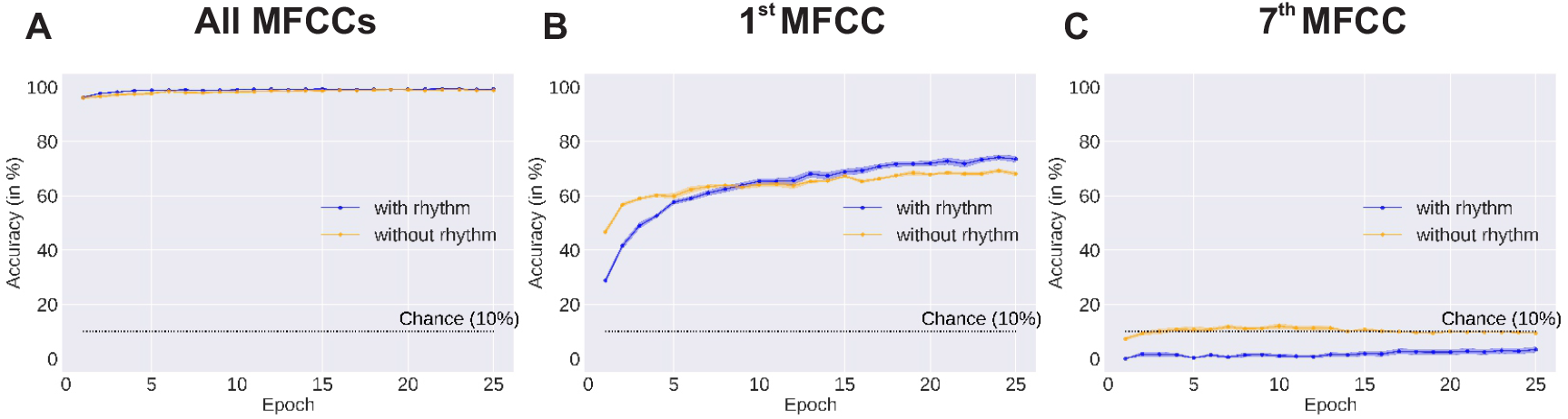
Performance with mel-frequency cepstral coefficients (MFCCs). Test accuracy per epoch of networks configured with frequency coefficients chosen uniformly at random from the range [1 Hz, 3 Hz] (blue). Values are averaged across ten random seeds. Shaded region depicts standard error. For reference, performance of the same network without endogenous oscillations is shown in yellow. Mel coefficients were **(A)** All, **(B)** 1st, **(C)** 7th.

**Figure S3:**
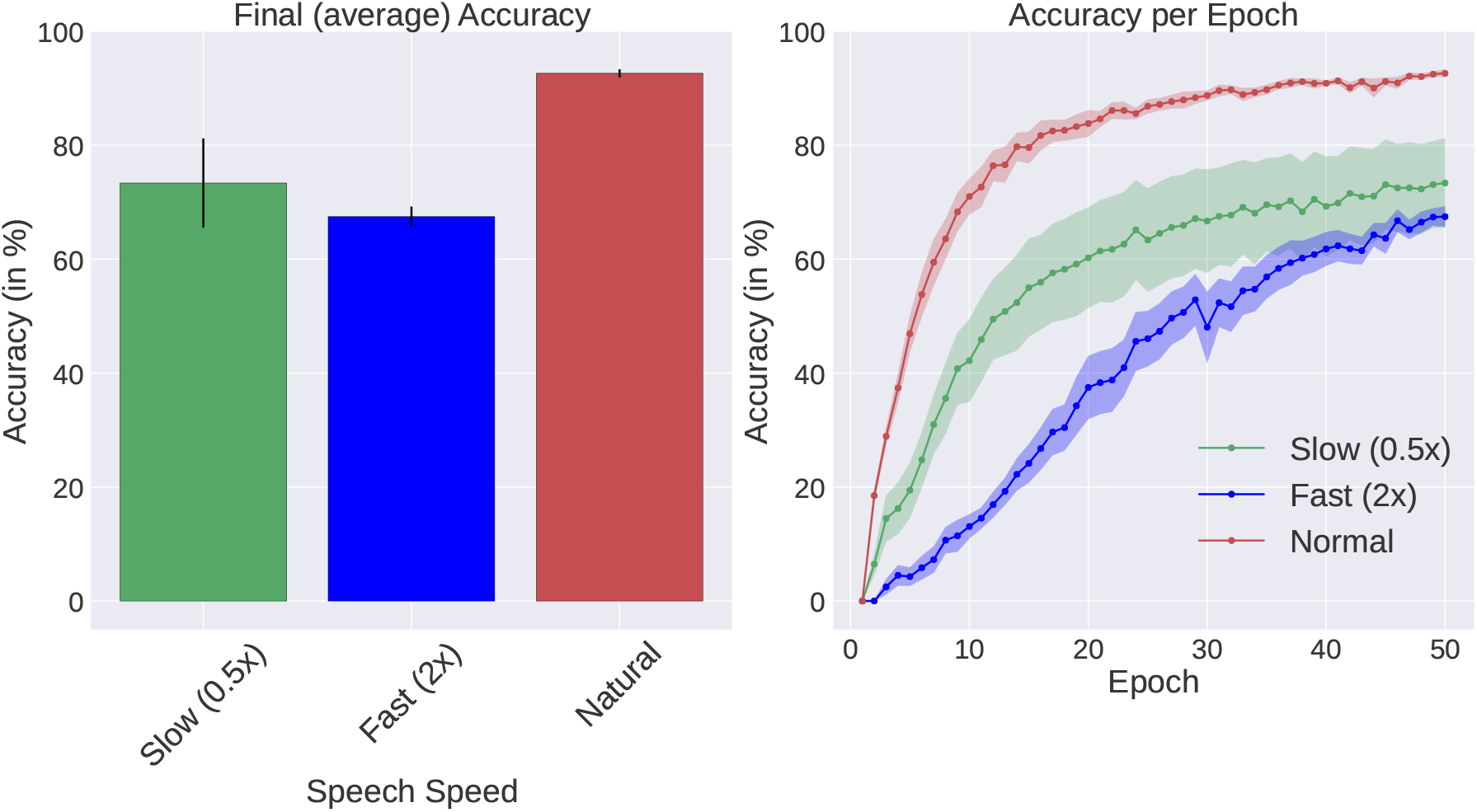
Effect of speech rate manipulation on performance. *Left:* Test accuracy of networks configured with [1 Hz, 5 Hz] frequency coefficients when the speech signal is slowed (0.5*x*), sped up (2*x*), and natural. *Right:* Test accuracy per epoch for the same networks. Values are averaged across ten random seeds. The shaded region depicts standard error.

**Figure S4:**
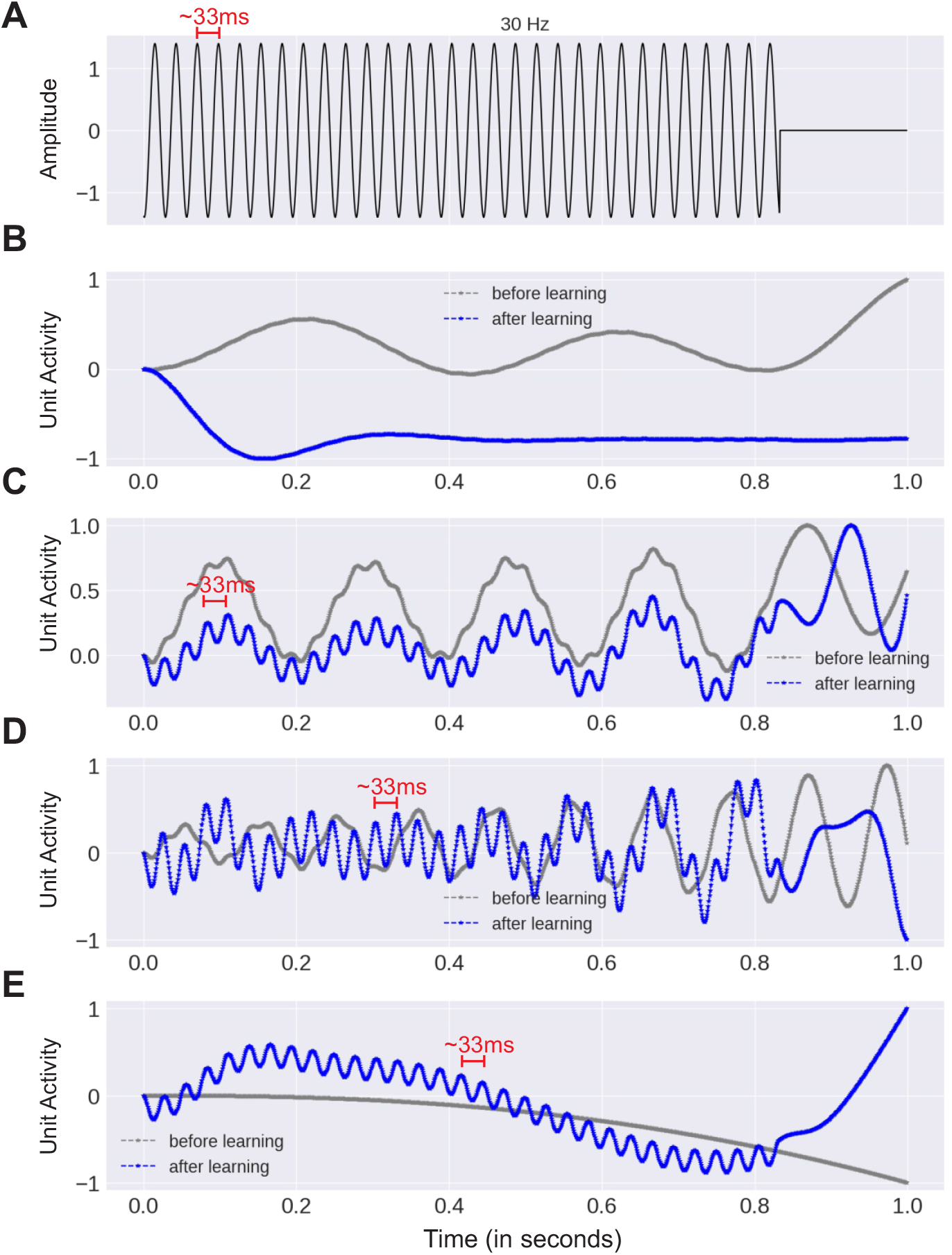
Network oscillations in response to a 30 Hz sinusoidal input. **(A)** Input waveform. **(B)** Response of an oscillator in a network configured with frequency coefficients drawn uniformly at random from the range [1 Hz, 10 Hz], **(C)** [10 Hz, 100 Hz], **(D)** [100 Hz, 110 Hz]. **(E)** Response when the frequency coefficients are set to 0. Oscillator activity is plotted as its normalized amplitude (scaled to [-1,1]). All networks, except for the network in **B**, tracks the temporal modulations of the input and correctly classifies the signal.

**Figure S5:**
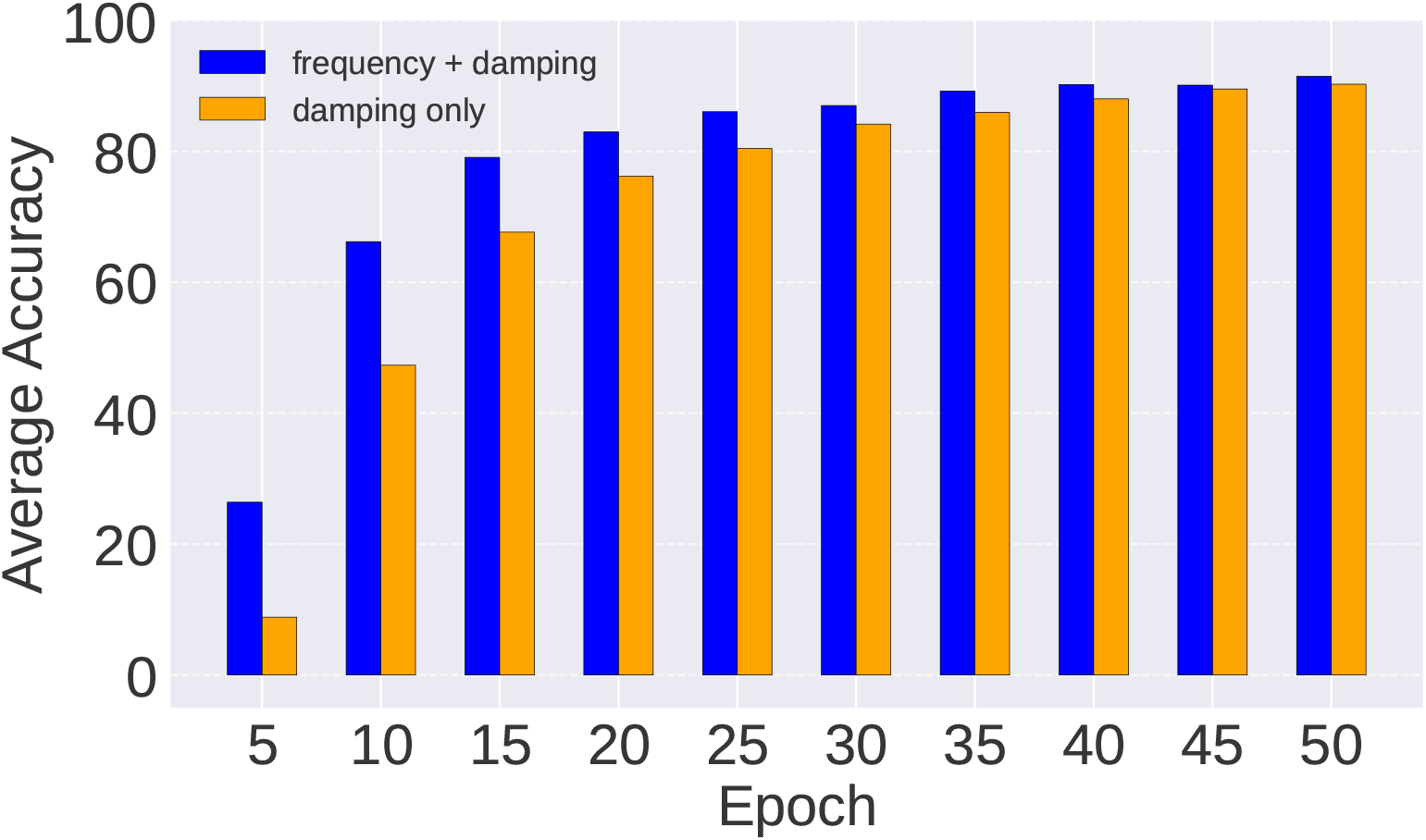
Task accuracy with damping only. Blue: Test accuracy on AudioMNIST (after every five epochs of training) of a network configured with frequency and damping coefficients drawn uniformly at random from the range [1 Hz, 3 Hz] and [0.1, 80], respectively. Yellow: same as blue, but with frequency coefficients set to 0. This network learns at a slower pace but eventually reaches high accuracy once it learns to oscillate with the speech envelope. Performances shown are averaged cross three random seeds.

**Figure S6:**
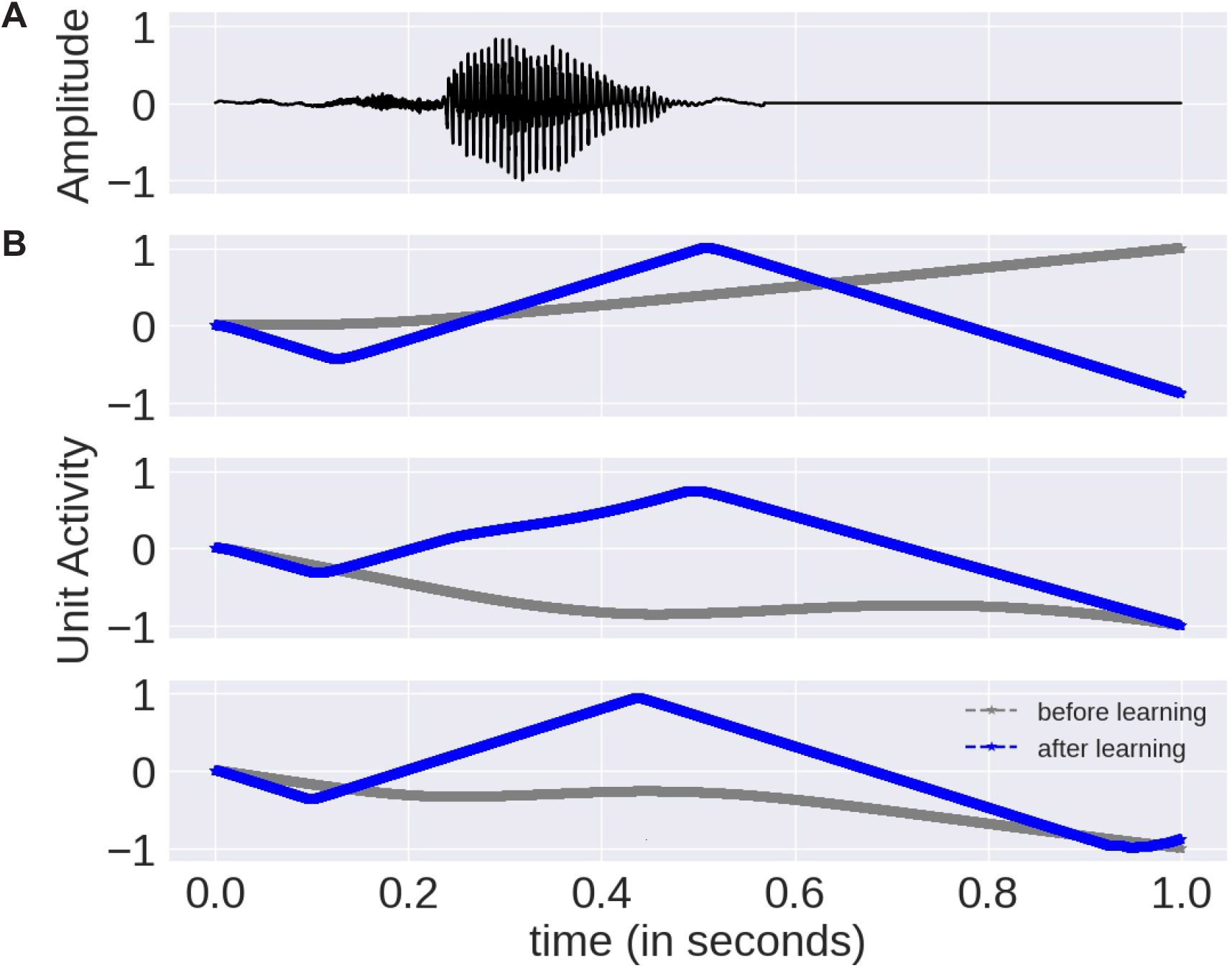
Oscillations in damped networks. **(A)** Waveform of the spoken word “three.” **(B)** Example response of three network oscillators before (grey) and after training (blue) on AudioMNIST. The frequency coefficient of the network is set to 0 and the damping coefficients are drawn uniformly at random from the range [0.1, 80]. Before training, no clear oscillatory pattern is present. After training, the oscillators exhibit rhythmic activity at approximately 1 Hz, matching the approximately 1 Hz rhythm of the speech envelope. The network classified the input correctly after training. Oscillator activity is plotted as its normalized amplitude (scaled to [-1, 1]).

**Figure S7:**
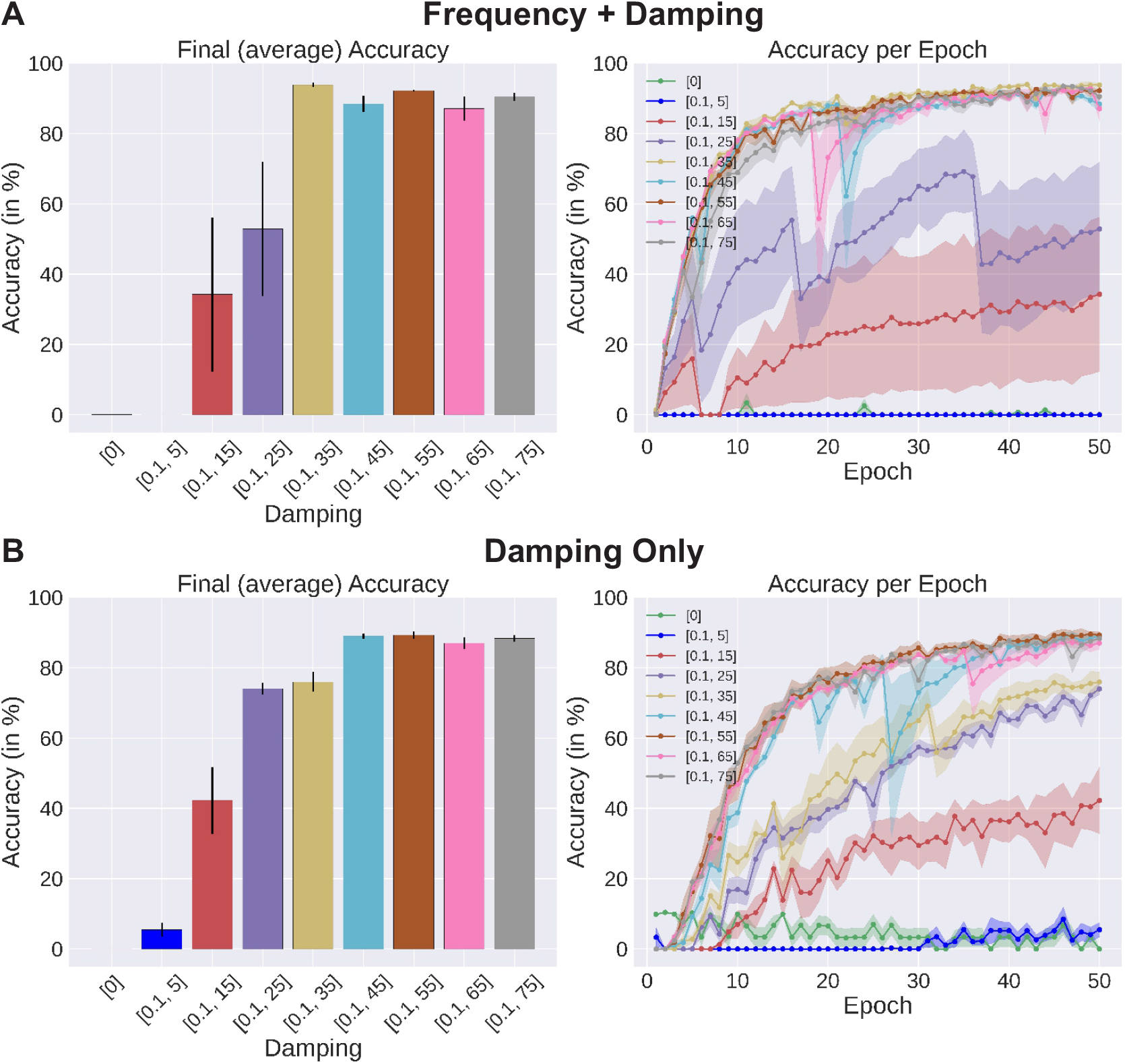
Impact of damping coefficient on network performance. **(A)** Accuracy in detecting English spoken words. Left: Test accuracy of networks configured with damping coefficients drawn uniformly at random from the specified ranges and oscillator frequencies drawn from the range [1, 3] Hz. Values are averaged across three random seeds. Error bars depict standard error. Right: Test accuracy after every epoch of training. **(B)** Performance of networks with the same damping coefficients but with the frequency coefficient set to 0, illustrating a similar dependence on damping.

**Figure S8:**
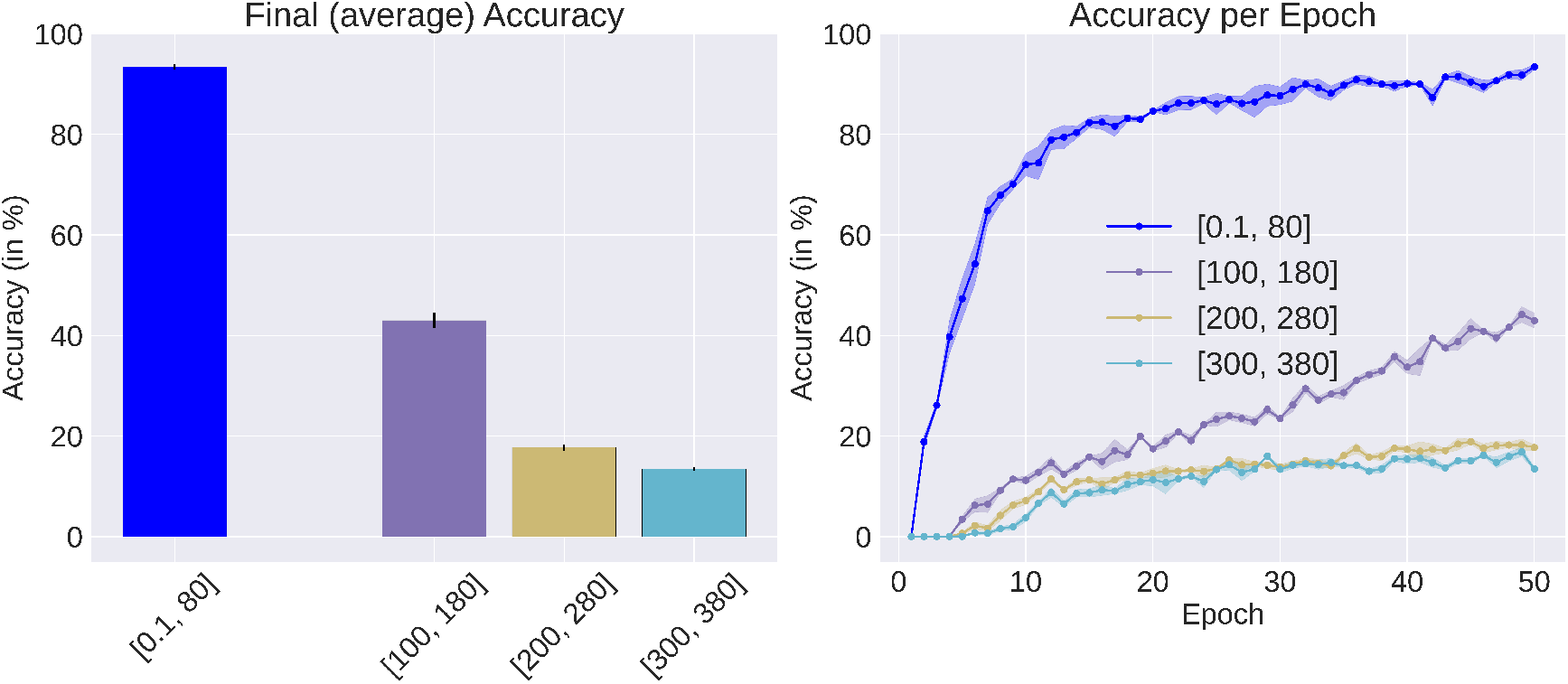
Impact of high damping coefficient on network performance. Accuracy in detecting English spoken words. Left: Test accuracy of networks configured with damping coefficients drawn uniformly at random from the specified ranges and oscillator frequencies drawn from the range [1, 3] Hz. Values are averaged across three random seeds. Error bars depict standard error. Right: Test accuracy after every epoch of training. Overdamping the network substantially reduces performance.

**Figure S9:**
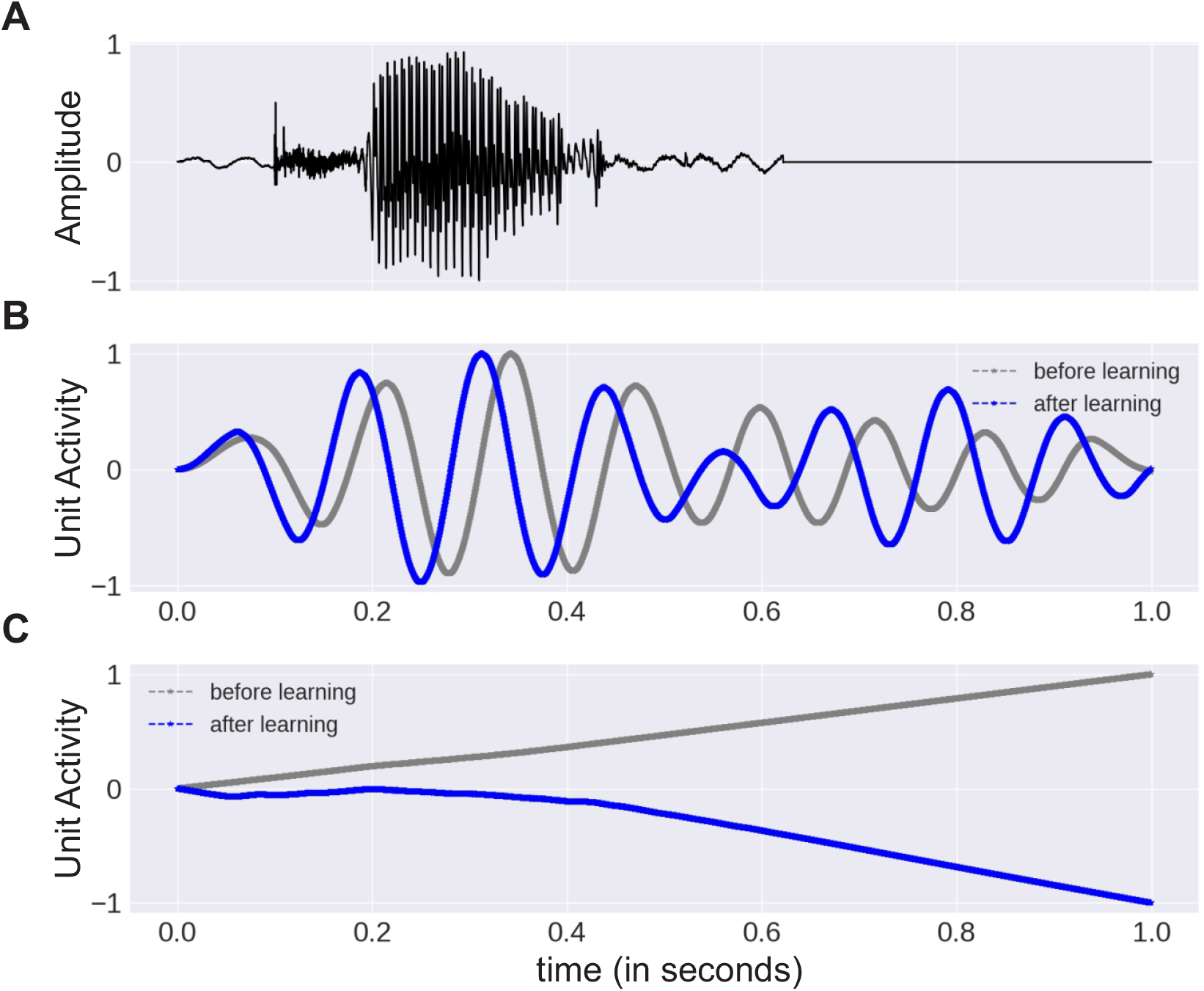
Network oscillations as a result of no damping and excessive damping. **(A)** Waveform of the spoken word “two” from the AudioMNIST dataset. **(B)** Example response of an oscillator in a network with damping coefficients set to 0 and frequency coefficients drawn from the range [1, 3] Hz. **(C)** Example response of an oscillator in a network with very high damping (*ϵ* = 240) and frequency coefficients drawn from the range [1 Hz, 3 Hz]. Oscillator activity is plotted as its normalized amplitude (scaled to [-1,1]). Both networks classified the input incorrectly after training on the full dataset.

